# Ecology not genetics explains correlated trait divergence during speciation

**DOI:** 10.1101/2024.07.02.601691

**Authors:** Clarissa F. de Carvalho, Nicholas P. Planidin, Romain Villoutreix, Víctor Soria-Carrasco, Rüdiger Riesch, Jeffrey L. Feder, Thomas L. Parchman, Jon Slate, Zachariah Gompert, Patrik Nosil

## Abstract

The formation of new species often involves the correlated divergence of multiple traits and genetic regions. However, the mechanisms by which such trait covariation builds up remain poorly understood. In this context, we consider two non-exclusive hypotheses. First, genetic covariance between traits can cause divergent selection on one trait to promote population divergence in correlated traits (a genetic covariation hypothesis). Second, correlated environmental pressures can generate selection on multiple traits, facilitating the evolution of trait complexes (an environmental covariation hypothesis). Here, we test these hypotheses using cryptic coloration (controlled by an incipient supergene) and chemical traits (*i.e.,* cuticular hydrocarbons, CHCs) involved in desiccation resistance and mate choice in *Timema cristinae* stick insects. We first demonstrate that population divergence in color-pattern is correlated with divergence in some (but not all) CHC traits. We show that when correlated population divergence does occur, it is unlikely to be explained by genetic covariation because within-population genetic covariance between color-pattern and CHCs traits is weak. In contrast, we find that correlated variation in climate and host plant likely generates selection jointly on color-pattern and some CHC traits. This supports the environmental covariation hypothesis, likely via the effects of two correlated environmental axes selecting on different traits. Finally, we provide evidence that misalignment between natural and sexual selection also contributes to patterns of correlated trait divergence. Our results shed light into transitions between phases of speciation by showing that environmental factors can promote population divergence in trait complexes, even without strong genetic covariance.

## Introduction

Species formation entails the progressive accumulation of phenotypic and genetic differences that generate reproductive isolation along a spectrum from weak to strong elimination of gene flow (1–3). This process unfolds over extended timescales that are usually impractical to observe in its entirety. Thus, researchers often investigate varying levels of divergence among existing populations and species as proxies for different stages of the speciation process (*i.e.,* the speciation continuum) (4–6). As such, by examining trait differences and reproductive isolation among populations, it may be possible to infer the processes underlying the transition from weak to strong population divergence (*i.e.,* different phases of the speciation process).

Numerous processes can contribute to population divergence along the speciation continuum. For instance, distinct environmental conditions can exert divergent selective pressures that diminish the homogenizing effects of gene flow. This process can lead to ecological isolation that fosters population divergence (7, 8). Given that local adaptation can be complex, numerous traits can be simultaneously under divergent selection. Thus, patterns of covariance will develop among these traits and their encoding genes as speciation proceeds (9, 10). This process can lead to alternate coadapted gene complexes, comprised of distinct sets of traits and genes that differentially adapt populations to contrasting selection pressures (11, 12), ultimately resulting in ecotypes and species that exhibit partial or strong reproductive isolation (13). From this perspective, a key challenge of speciation studies is to elucidate the mechanisms that enable the adaptive divergence of traits and genes into differentially correlated complexes, the entities we recognize as ecotypes and species. We here investigate the mechanisms by which divergence in adaptive traits can become correlated (*‘*correlated trait divergence’, hereafter, Fig. 1).

**Figure 1.**
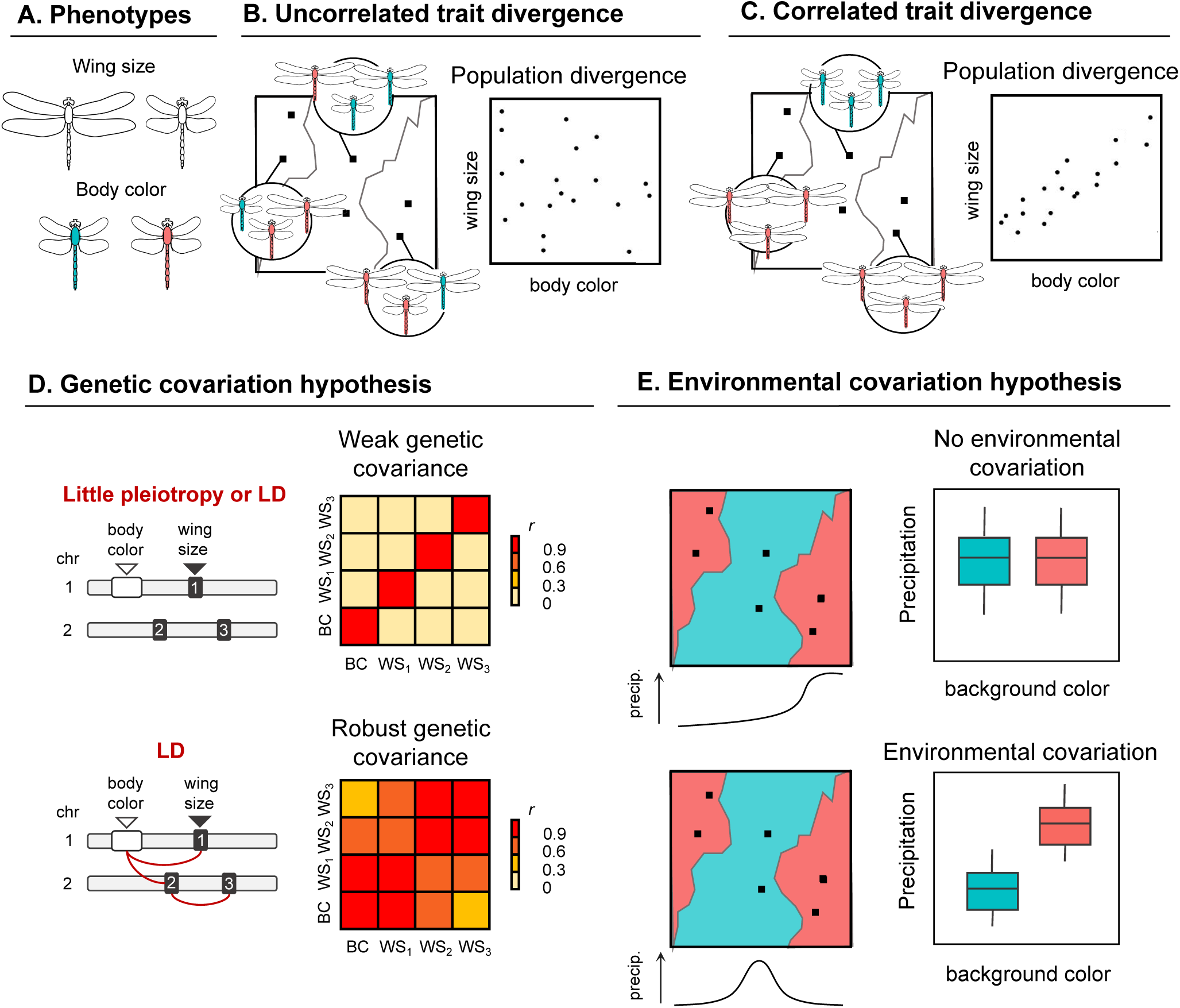
Hypotheses to explain correlated trait divergence. (**A**) Phenotypes for this hypothetical example. (**B**) Example of when population divergence in wing and in body color are not correlated. (**C**) Example of correlated trait divergence among populations. Greater divergence in wing size is correlated with greater divergence in body color. Two general and not mutually exclusive hypotheses are proposed to explain correlated trait divergence. (**D**) The ‘genetic covariation’ hypothesis predicts that genetic covariance between traits leads to correlated trait divergence. In this example, linkage disequilibrium (LD) between the gene controlling body color (BC) and the three genes controlling wing size (WS1-3) generates positive genetic covariance. (**E**) The ‘environmental covariation’ hypothesis predicts that correlated selective pressures lead to correlated trait divergence. In this example, background color influences body color and precipitation levels (precip.) influence wing size.

Extensive research spanning diverse taxa has demonstrated examples of correlated trait divergence. For example, studies in fish (14–17), mammals (18), mollusks (19), insects (20, 21), and plants (22–24) have revealed correlated divergence between traits. Despite these findings, the mechanisms by which divergence in different adaptive traits becomes correlated is less understood. In addition, to which extent these correlated trait complexes represent phenotypic polymorphisms (*i.e.*, morphs) controlled by simple genetics (*e.g.*, major loci or chromosomal inversions) or divergent ecotypes with some degree of genome-wide differentiation and reproductive isolation remains unclear. Consequently, the transition between morphs to ecotypes and species requires further study. This gap in our knowledge can be investigated by analyzing divergence in various traits across a continuum of divergence.

We provide such a study here, based on wild stick insect populations spanning a wide range of divergence in adaptive traits and reproductive isolation. We consider two non-mutually exclusive hypotheses to test for and explain correlated trait divergence (Fig. 1). The first hypothesis posits that the genetic covariance of traits causes their correlated divergence (*i.e.,* a genetic covariation hypothesis; Fig. 1D). This is because genetic covariance arising from pleiotropy or linkage disequilibrium (*i.e.,* LD) can result in indirect correlated responses among traits, where selection acting on one trait impacts the response of other traits (25, 26). This way, divergence among populations tend to occur along axes with greater genetic variation and covariation (27). Thus, if the genetic covariation hypothesis holds true, we predict significant genetic covariance between locally adapted traits, which if due to physical linkage can also represent supergene evolution (18–20, 24, 28).

The second hypothesis posits that the environmental variation across the landscape generates correlated selection on different traits, resulting in their correlated divergence (*i.e.,* an environmental covariation hypothesis)(Fig. 1E). This phenomenon can manifest itself in two ways. First, when a single selective pressure influences multiple traits concurrently, such as when high predation impacts multiple characters in guppies (29). Second, when environmental gradients are structured across the landscape in a manner that causes multiple environmental pressures covary. The correlated selective pressures arising from this covariation can thus jointly affect traits with no genetic or functional relationship with each other (30). For example, higher predation intensity in guppy populations is often correlated with other abiotic factors such as higher temperatures and light intensity, which affect multiple traits in guppies (29). In any case, if the environmental covariation hypothesis holds true, we predict that the environmental factors exerting selective pressures on different traits would be correlated with each other and with trait co-divergence. This process could be further influenced by the alignment between different selective forces acting on the traits, such as a different source of natural selection or sexual selection (31–34).

We investigate correlated trait divergence and test these two hypotheses using *Timema cristinae* stick insects, an emerging model system for studying adaptation and speciation. These wingless, plant-feeding insects are distributed throughout the Santa Ynez mountains in California, USA (35). *Timema cristinae* is primarily found in two host-plant species: *Adenostoma fasciculatum* (Rosaceae), and *Ceanothus spinosus* (Rhamnaceae) (36, 37). Divergent selection associated with the two host-plant species contributes to partial reproductive isolation between populations associated to them, leading to the *Adenostoma* and *Ceanothus* ecotypes (38). Here, we specifically investigate two types of adaptive traits that vary between the ecotypes (Fig. 2): (1) cryptic color-pattern polymorphism; and (2) cuticular hydrocarbons (CHCs, hereafter).

**Figure 2.**
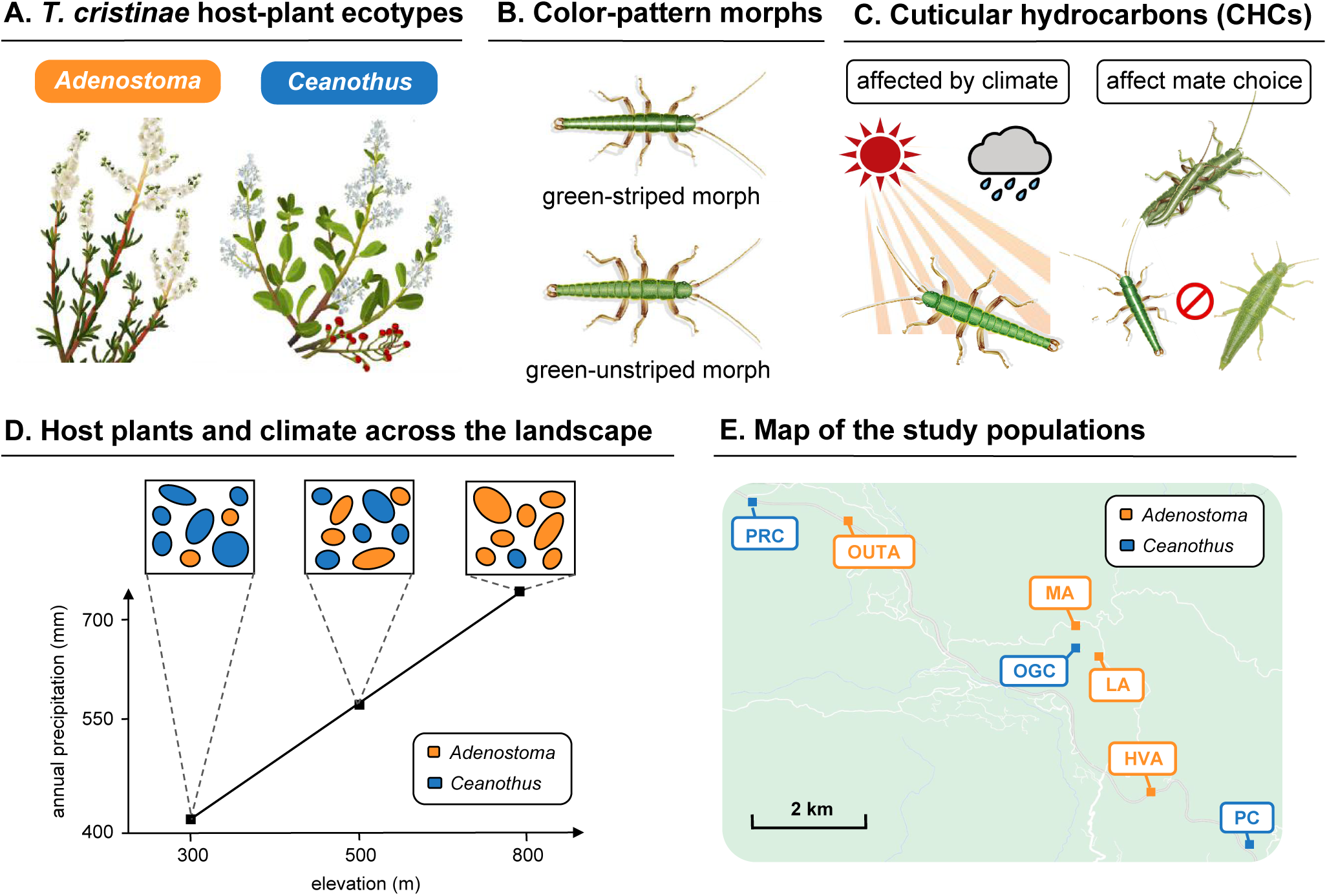
*Timema cristinae* ecotypes and the traits investigated in this study. (**A**) Host-plant species which define the ecotypes. (**B**) Color-pattern morphs in *T. cristinae*, mainly selected by host-plant species. (**C**) Female cuticular hydrocarbons (CHCs), with functions in chemical communication and water balance. Cuticular hydrocarbons are associated with climatic adaptation and they affect mate choice, thus are also influenced by sexual selection. (**D**) Abundance of host-plant species and climatic variables tend to vary with elevation across the landscape where *T. cristinae* is found. (**E**) Map of the populations used in this study. The association between host-plant species and climate was taken into account in the selection of study sites (*i.e.*, populations on *Adenostoma* and *Ceanothus* across different elevations).

Cryptic color-pattern morphs in *T. cristinae* are characterized by the presence versus absence of a white dorsal stripe. This trait is subject to divergent selection exerted by visual predators such as birds and lizards (36, 39). The green-striped morph is more cryptic on the needle-like leaves of *Adenostoma* plants, while the green-unstriped morph is more cryptic on the broader leaves of *Ceanothus* (Fig. 2). Accordingly, the frequencies of the color-pattern morphs vary between host-plant ecotypes, with the striped morph being more prevalent on the *Adenostoma* ecotype and the unstriped morph predominating on the *Ceanothus* ecotype (36, 39–41). Although *T. cristinae* has a third morph that is melanistic (dark) in body color and lacks a stripe (42), its frequencies do not exhibit significant host-related divergence between populations (43). Therefore, it is not the focus of this study. Despite the pronounced selection acting on the color-pattern morphs, a combination of gene flow and negative frequency-dependent selection prevents fixation of either the green-striped or green-unstriped morph in the populations, creating a geographic mosaic in the degree of color-pattern divergence among populations (41, 44). Recent research has revealed that color-pattern is strongly associated with two large regions of suppressed recombination on chromosome 8, referred to as the *Pattern* and *Mel-Stripe2* loci. (44). Together, these two loci thus function as an incipient supergene (see (44)).

In addition to color-pattern, we examine CHCs, which are waxy chemical compounds with roles in adaptation, sexual selection, and speciation in insects. For example, CHCs have been shown to contribute to maintenance of the water balance, communication, and mate recognition in insects (45, 46). The functions played by the CHCs vary according to some properties of these compounds, such as their carbon chain length (46). In *T. cristinae*, it has been shown that CHCs are controlled by multiple loci across the genome, exhibiting non-zero but modest heritability (47, 48). A recent study has further reported that the single nucleotide polymorphisms (SNPs) linked to CHC variation overlap with genetic regions associated with climate adaptation in different *Timema* species (48). This link between CHCs and climate adaptation is most likely related to their general role in maintaining water balance and preventing desiccation (49–51). It has also been shown that CHC profiles differ between the *T. cristinae* host-plant ecotypes (47), but the extent to which this relationship reflects adaptation remains unclear. Most critically, greater divergence in female CHCs in *Timema* is associated with mate choice and sexual isolation, both within and between species (47, 52). This was demonstrated by manipulative perfuming experiments, which have established a causal role for female CHCs in male mate choice within a population of *T. cristinae* and sexual isolation between a species pair (47). These results prompt our focus on female rather than male CHCs in the current manuscript.

Previous research in *T. cristinae* has shown that the effects of color-pattern divergence on genetic differentiation are restricted to chromosome 8, while divergence in female CHC is associated with greater genome-wide differentiation (47). However, these patterns were analyzed independently, leaving unanswered questions regarding the potential cumulative effects of differentiation in each trait, the extent to which they show correlated trait divergence, and why. Therefore we seek to bridge this gap here by exploring correlated trait divergence between color-pattern and female CHCs among populations.

In our study, we first test whether color-pattern and female CHC traits show correlated divergence among populations spanning different stages of divergence. We subsequently test the genetic covariation hypothesis by testing the prediction that there is within-population genetic covariance between color-pattern and CHCs. We then test the environmental covariation hypothesis by exploring the relationship between the environmental conditions that exert selection on color-pattern (*e.g.*, host plants) and on chemical traits (*e.g.*, climate) across *T. cristinae* populations.

Specifically, we test the prediction that these environmental axes are correlated. Given the importance of CHCs in sexual isolation, we take advantage of abundant experimental data on mate choice in *T. cristinae* to also explore whether sexual selection may affect patterns of correlated divergence between color-pattern and CHCs. Finally, we integrate our results with existing knowledge on differentiation and reproductive isolation among *Timema* species, generating broader understanding of transitions between phases of speciation.

## Results

### Population divergence in color-pattern and in CHCs

We begin our investigation by estimating population divergence in color-pattern and CHCs. We do so by using phenotypic data from (47), encompassing seven populations of *T. cristinae* at different stages of divergence (*i.e.*, comprising 21 population pairs when population divergence was considered, Table S1). We assess divergence in color-pattern among populations based on the frequency of the striped morph in the population (*i.e.,* green-striped versus green-unstriped). Divergence in female CHC traits is evaluated by considering three classes of molecules classified by their carbon chain length: pentacosanes, heptacosanes, and nonacosanes (*i.e.*, chains of 25, 27, and 29 hydrocarbons, respectively). In this work, we analyze each CHC class as a separate trait (see Material and Methods). We find a broad spectrum of diversity across all traits. For instance, with respect to color-pattern, the striped morph frequency within populations range from as low as 1% (PRC) to as high as 86% (LA). We thus move on to test the co-divergence between color-pattern and the different CHC traits.

### Divergence in color-pattern is correlated with divergence in some CHC traits

We find that patterns of co-divergence with color-pattern differed among CHC traits. Our analyses indicate that population divergence in color-pattern is strongly correlated with divergence in nonacaosanes (*r*=0.82, *P*<0.001, Mantel test; Fig. 3), and is modestly correlated with divergence in heptacosanes (*r*=0.38, *P*=0.10, Mantel test). However, color-pattern divergence among populations exhibits no significant correlation with female pentacosanes (*r*=0.11, *P*=0.27, Mantel test; Fig. 3). We thus next test hypotheses for the causes of this variation.

**Figure 3.**
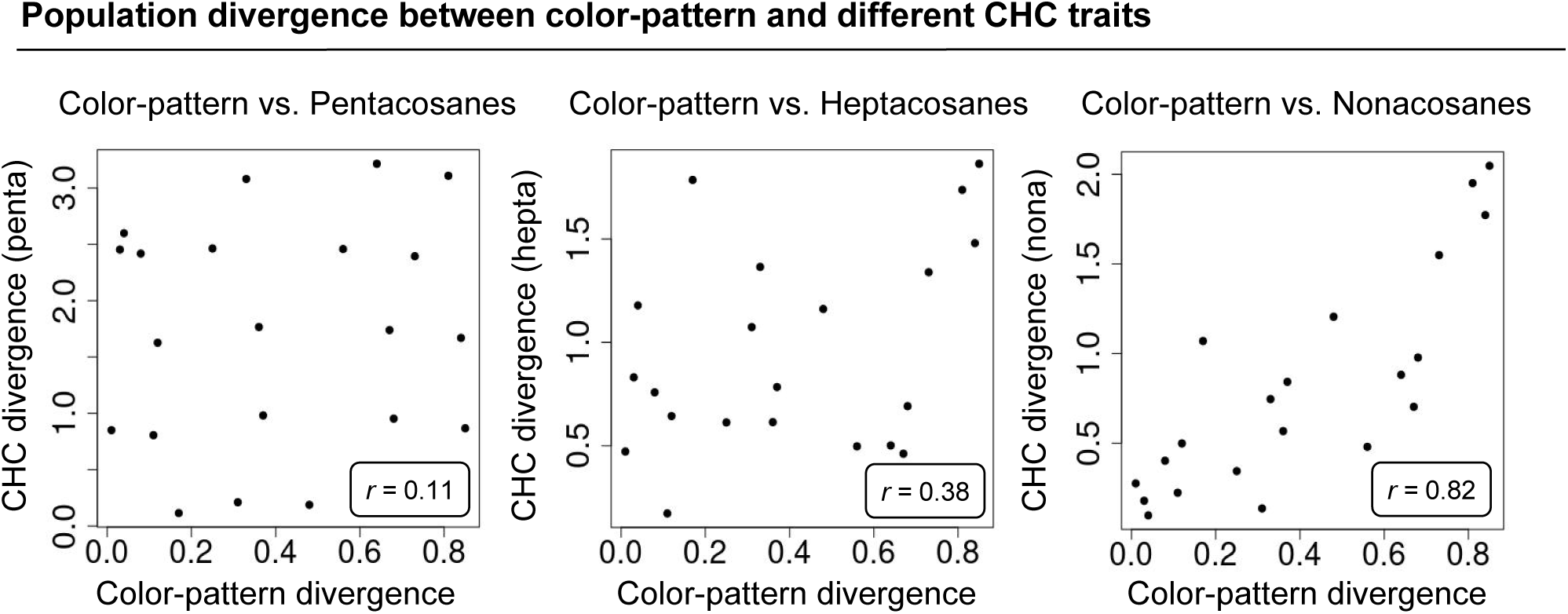
Correlated divergence between color-pattern and female CHC traits. Population divergence between color-pattern and pentacosanes (*r*=0.11, *P*=0.27), heptacosanes (*r*=0.38, *P*=0.10), and nonacosanes (*r*=0.82, *P*<0.001, Mantel tests). Among the CHC traits, nonacosanes are the class with strongest correlated divergence with color-pattern.

### Weak genetic covariance between color-pattern and CHCs within the same population

We test the genetic covariation hypothesis by quantifying the genetic covariance between color-pattern and each of the CHC traits within the population FHA (Far Hill, *Adenostoma*, 34.518 N, - 119.801 W). Our results show modest genetic covariance values (at best) between color-pattern and all CHC traits, being *r* = 0.10 for nonacosanes (Pearson correlation, 95% CIs = −0.04 – 0.24, Fig. 4A), *r* = −0.06 for heptacosanes (Pearson correlation, 95% CIs = −0.20 – 0.09), and *r* = −0.16 for pentacosanes (Pearson correlation, 95% CI = −0.32 – 0.00, Fig. 4A). Thus, our findings do not provide strong support for the genetic covariation hypothesis, urging a test of the environmental covariation hypothesis.

**Figure 4.**
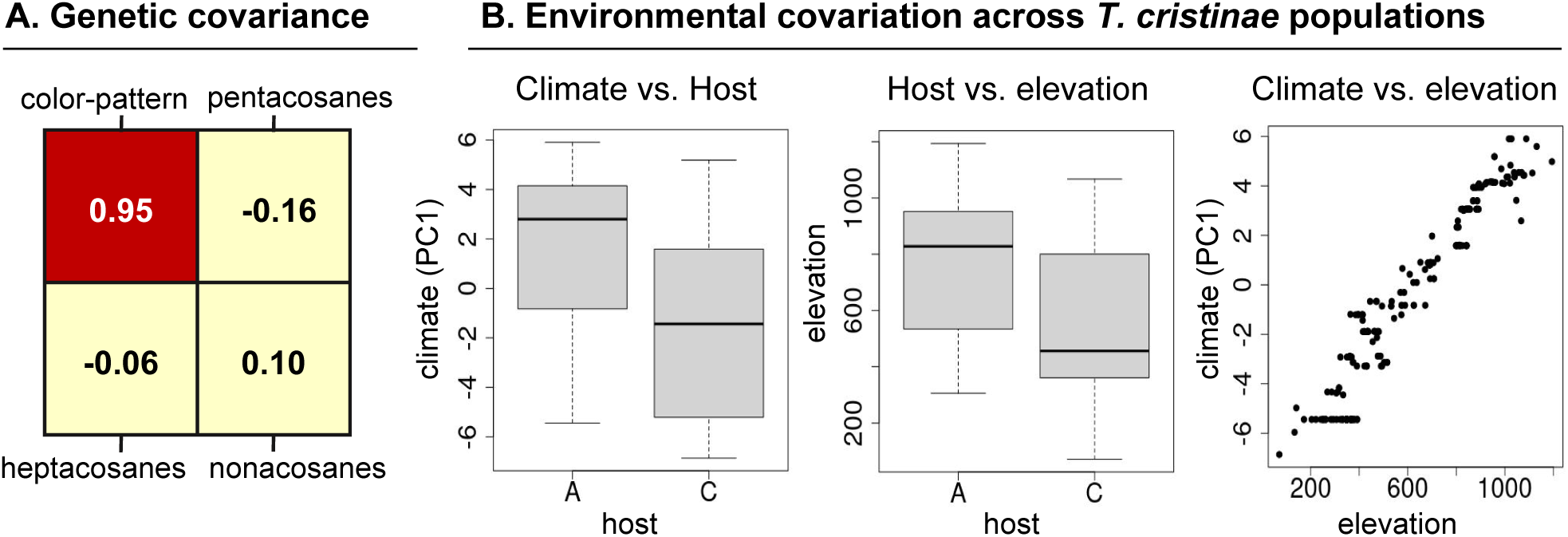
Processes explaining correlated trait divergence between color-pattern and CHC traits, especially nonacosanes. (**A**) Genetic covariance between color pattern and the female CHC traits (yellow), represented by Pearson correlation coefficients. Based on these results, genetic covariance is unlikely to explain correlated trait divergence between color-pattern CHC traits. Heritability in color-pattern (the diagonal of the correlation matrix) is represented in red. (**B**) Environmental axes influencing color-pattern (host-plant species) and CHCs (climate). Host-plant species are strongly influenced by climate, represented by the first principal component axis summarizing 19 WorldClim variables (PC1=67% of variance explained; *W* = 7872, *P* < 0.001, Wilcoxon signed-rank test). This relationship is explained by the common association between each environmental variable and elevation (host-plant species and elevation; *W* = 7932, *P* < 0.01, Wilcoxon signed-rank test; PC1 summarizing climatic variables and elevation; *r* = 0.96, *P* < 0.001, Pearson correlation). These correlated selective pressures can explain correlated trait divergence. Abbreviations: A= *Adenostoma*, C=*Ceanothus*.

### Host plant and climate are associated with correlated trait divergence

To evaluate the environmental covariation hypothesis, we test the prediction that there is a correlation between the primary axes of selection for color-pattern versus CHC traits, *i.e.,* between host-plant species and climate (see Figs. S1-S2 for association between climate and CHCs). To test this prediction, we use data from (41), encompassing information from 206 *T. cristinae* populations, including details on host plant (*i.e.*, whether the population is of *Adenostoma* or *Ceanothus* ecotype) and climatic variables. We find a robust association between host-plant species and nearly all 19 bioclimatic variables from WorldClim (53), with the exception being the ‘maximum temperature of the warmest month’ (Table S3). This association is exemplified by the highly significnat correlation between host-plant species and the first principal component axis summarizing the climatic variables (*W* = 7872, *P* < 0.001, Wilcoxon signed-rank test; Fig. 4B). We find that this relationship likely stems from the shared variability of climate and host plant species with elevation (*r* = 0.96, *P* < 0.001, Pearson correlation; *W* = 7932, *P* < 0.001, Wilcoxon signed-rank test; respectively). This alignment of environmental pressures between climatic variables and host plants can explain the positive correlation observed between divergence in color-pattern with divergence in nonacosanes, and perhaps heptacosanes. In contrast, this does not appear to be the case for pentacosanes. We explore this topic below in the next section.

### Sexual selection explains weak correlated trait divergence

We consider another hypothesis for correlated trait divergence that explores the potential influence of sexual selection. We speciifically ask if sexual selection or mate choice based on CHCs may explain the weak correlation between population divergence in pentacosanes and in color-pattern. Given the known role of female CHCs in mate choice and sexual isolation in *Timema* (47), it is plausible that sexual selection is shaping the divergence of CHCs across populations. Natural and sexual selection pressures could thus be simultaneously driving variation in female CHCs in different ways for different CHC traits (31, 33, 34). Therefore, we posit a scenario where a potential misalignment between natural and sexual selection pressures could disrupt the environmental covariation from driving correlated trait divergence of traits (Fig. 5). If this hypothesis holds true, we predict that the divergence in pentacosanes should exhibit a strong correlation with sexual isolation levels among populations (rather than with color-pattern, as reported above).

**Figure 5.**
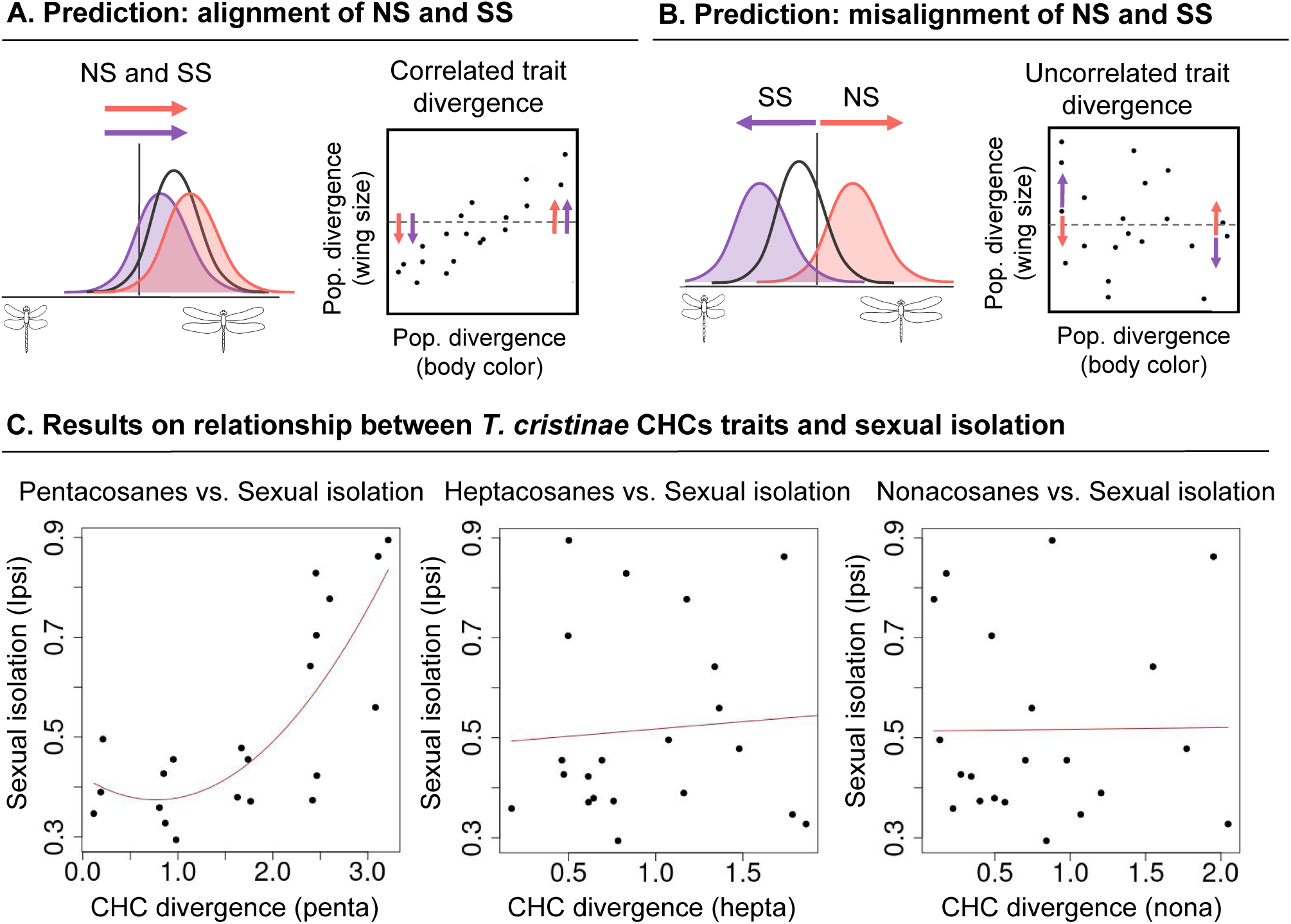
Hypothesis of misalignment between natural (NS) and sexual (SS) selection to explain lack of correlation in trait divergence. In this example, wing size is influenced by NS and SS (pink and purple, respectively). When these two pressures are aligned (**A**), correlated trait divergence can occur between wing size and body color. However, when NS and SS are misaligned (**B**), the phenotypic means can be shifted away from natural selection’s optimal adaptive peak. Consequently, SS disrupts the correlated NS effects in wing size and body color, resulting in their divergence to not be correlated. (**C**) Association between the different CHC traits and sexual isolation. Divergence in pentacosanes shows strong association with sexual isolation (Bayesian linear mixed models, BLMM, *β*=0.76 [0.49, 1.03 95% ETPI]), suggesting a role for sexual selection. Meanwhile, divergence in heptacosanes (*β*=0.09 [−0.38, 0.56 95% ETPI]) and nonacosanes (*β*=0.02 [−0.45, 0.48 95% ETPI]) are not significantly correlated with sexual isolation (see Table S4 for best models’ fit in BLMM). The strong correlation between pentacosane divergence and sexual isolation suggests that a misalignment between SS and NS (exerted by climate) could be disrupting the environmental covariation, thus resulting in uncorrelated divergence between pentacosanes andcolor-pattern. Abbreviations: pop. = population.

To test this prediction, we estimated correlations between divergence in different female CHC traits and the pairwise index for sexual isolation (*I_PSI_*) among populations using the data from Riesch *et al.* (47). We find a strong positive relationship between divergence in pentacosanes and sexual isolation (Bayesian linear mixed models, BLMM, *β*=0.76, 95% equal tail probability interval, ETPI = 0.49 – 1.03, a quadratic model; Fig. 5B; Table S4). Furthermore, BLMM reveal that divergence in female pentacosanes explain sexual isolation slightly better than geographical distance (Table S5). In contrast, heptacosanes and nonacosanes did not exhibit significant associations with sexual isolation (BLMM *β*=0.09, 95% ETPI = −0.38 – 0.56; *β*=0.01, 95% ETPI = −0.45 – 0.48; respectively). Thus, sexual selection could be exerting effects mainly on pentacosanes, potentially in misalignment with the effects of natural selection deriving from climate.

## Discussion

Speciation is often a complex process involving the divergence of various genetic loci and traits. When multiple traits co-diverge, their synergistic interaction can lead to enhanced phenotypic and genetic differentiation, potentially strengthening isolating barriers between populations. Therefore, understanding correlated divergence in adaptive traits and the mechanisms underlying it can be crucial for understanding the processes by which population divergence can build up through different phases of speciation. In this study, we demonstrate that the correlated divergence of adaptive traits in *T. cristinae* is driven more by environmental covariation than by genetic covariation. Furthermore, our findings suggest that the effects of sexual selection contrast the effects of natural selection, affecting correlated trait divergence. Our analyses and results on the mechanisms that underpin correlated trait divergence contribute to knowledge concerning transitions between phases of speciation (54, 55). Discussion of our main findings follows.

### Environmental covariation drive trait co-divergence, despite weak genetic covariation

In this study, we find significant correlated divergence between color-pattern and one class of female CHCs (*i.e.,* nonacosanes, and, to lesser extent, heptacosanes). However, we find weak genetic covariance between color-pattern and all CHC traits analyzed within the FHA population, challenging the hypothesis of a strong genetic link between these traits, and consistent with the fact that these two types of traits are governed by distinct genetic architectures in *T. cristinae*. While color-pattern is under the control of an incipient supergene with high heritability (41, 42), CHCs are influenced by multiple genetic regions with modest heritability (47, 48). Thus, there do not appear to be shared loci of large effect pleiotropically influencing both traits (*i.e.*, loci with effects on CHCs do not appear to be clustered or preferentially located within the supergene region governing color-pattern). Consequently, our study implies that selection acting on one of these traits may not necessarily drive evolutionary changes in the other, indicating that their genetic covariance is unlikely to contribute to their correlated divergence.

Instead, our results show that environmental covariation across the landscape where *T. cristinae* lives can better explain the correlated divergence observed between color-pattern and CHC traits. Our results reveal an alignment of the environmental axes that exert correlated selection on each trait. Specifically, host-plant species and climatic variables strongly correlate in space due to their association with elevation. For color-pattern, this relationship reflects a difference in the abundances of hosts with elevation, while for the CHC traits nonacosanes and heptacosanes it may reflect differences in temperature and precipitation. In this regard, nonacosanes, whose divergence showed the strongest covariance with color-pattern, contain longest carbon chains. Longer chained CHCs tend to have higher melting temperatures and, as a result, are associated with a greater ability to maintain the insects’ water balance (46, 56). Thus, the findings for nonacosanes are consistent with a role for the these CHCs in creating a water-proof layer that varies in an elevation-dependent manner. Therefore, our results align with the concept of environmental covariation generating correlated trait divergence, even in the absence of strong genetic covariance. Our results challenge a prevailing view in the literature that underscores genetic and developmental causes for trait covariation compared to the environment (29, 30). Nonetheless, the mixed support and role for genetic covariance in adaptive divergence has been highlighted in a past meta-analysis and our results are consistent with this past work (57). Furthermore, our findings refine current emphasis on the role of specific genetic architectures that suppress recombination in evolution, implying that such architectures can facilitate adaptation and supergene evolution without necessarily contributing to genome-wide differentiation and speciation (28, 58).

### Contrasting effects of natural and sexual selection

Our results also reveal that divergence in pentacosanes is not correlated with divergence in color-pattern. That is, although pentacosanes are influenced by climatic conditions and are likely subjected to their natural selection pressures (Figs. S1-S2)(48), the environmental covariation between climate and host-plants seems to be insufficient to generate correlated divergence between pentacosanes and color-pattern. Instead, divergence in pentacosanes is more correlated to sexual isolation among populations. This association is quadratic, potentially indicating the nonlinear impacts of trait evolution on mate choice and reproductive barriers (59–61). That is, our results show a rapid evolution of sexual isolation once populations reach threshold levels of divergence in female pentacosanes. Conversely, neither heptacosanes nor nonacosanes show significant association with sexual isolation, suggesting that the results found in (47) were mainly linked to variation in pentacosanes.

Collectively, our findings imply that the interplay between natural and sexual selection may lead to contrasting evolutionary trajectories for pentacosanes. While natural selection may drive the optimization of CHCs for climatic adaptation, sexual selection may favor specific chemical profiles for successful mate attraction and recognition (62). This scenario has been previously described in crickets, whose CHC profiles contrasted significantly under the effects of natural versus sexual selection (63), and may occur more generally for traits that experience natural and sexual selection (31–34). As such, our results suggest that, although sexual selection on *T. cristinae* pentacosanes contributes to the evolution of sexual isolation among populations, it may disrupt the correlated selective pressures that lead to correlated trait divergence with color-pattern. Our results align with meta-analyses indicating that sexual selection (mating selection) may exert stronger effects than viability selection (survival) (61). To further elucidate this phenomenon, future research in the system should quantify the strength and form of natural and sexual selection targeting different classes of female CHCs.

### Evolution from morphs to ecotypes to species

This study provides insights into the processes that promote the transition from morphs to ecotypes with variable levels of population differentiation to the more divergent units considered species. In terms of ecotypes, recent studies in *Timema* have shown how fluctuation can occur around very different equilibria, resulting in ‘balanced ecotypes’ that represent elements of polymorphism but also intermediate stages of ecological speciation (41, 44). Here, we build upon past work by showing a pivotal role for environmental covariation in driving correlated and more pronounced population divergence in *T. cristinae*. Despite this divergence, reproductive isolation among these populations is still far from complete (38). In this context, why has divergence of *T. cristinae* populations not progressed further towards speciation?

To shed some light on this question, we consider features that characterize distinct *Timema* species. Here, we highlight two key phenomena: genome-wide differentiation and strong divergence in CHCs (Fig. S3) (47, 52). These two factors may be required to create new *Timema* species, likely facilitated by periods of geographic isolation. In contrast, notable differences in body color or color-pattern do not always occur between *Timema* species and thus do not appear essential for species divergence. As such, we speculate that if correlated trait divergence does play a role in the later stages of speciation, it is more likely to involve traits other than body color or color-pattern, such as CHCs and intrinsic post-mating isolation. Therefore, the path to greater population divergence may sometimes lead to more complete speciation, as observed in *Rhagoletis* flies (64), but in other cases as reported here, it may not. We speculate that the outcome depends on the traits involved and their effect on adaptation versus reproductive isolation. Further studies are now required to test these hypotheses, using an even wider range of the divergence process such that the entire path to new species can be reconstructed along the speciation continuum.

### Conclusion

Here, we report correlated divergence between color-pattern morph variation and certain CHC traits, primarily attributed to environmental covariation across the landscape rather than to strong genetic covariance. Our results thus underscore a pivotal, but often understudied, role for environmental covariation in the speciation process. Interestingly, environmental variation at a smaller scale, that is the colors of leaves versus stems within plant individuals, has been implicated in the evolution of color morphs in the *Timema* system (65), opening the potential for further work to even more fully investigate the evolution of morphs to ecotypes to species. Indeed, the weak correlation between color-pattern and other CHC traits provided an opportunity to explore alternative explanations, shedding light on how a misalignment between sexual and natural selection might affect trait divergence and speciation. Our study highlights the importance of investigating multiple dimensions that drive population divergence, offering valuable insights into the processes that can facilitate or limit divergence and transitions between phases along the speciation continuum. Comparable joint tests of the roles of genetics and environment in other taxa will likely continue to shed light into the general processes driving and constraining speciation.

## Material and Methods

### Association between divergence in color-pattern and CHCs

We tested whether divergence in color-pattern is correlated with divergence in different CHC classes using data from Riesch *et al.* (47) (https://dx.doi.org/10.5061/dryad.nq67q). For color-pattern, we computed the Euclidean distances from percentage of striped individuals from seven populations (PC, HVA, MA, LA, OUTA, PRC, OGC; *i.e.,* comprising 21 population pairs when population divergence was considered, Table S1). For CHCs, we used data of females from the same populations. There were 25 compounds in total, summarized according to their chain length as pentacosanes, heptacosanes and nonacosanes (*i.e.,* 25, 27, and 29 carbons, respectively). We conducted principal component analyses (PCA) separately for each CHC class, based on a covariance matrix with promax rotation. We then retained the principal component axes with an eigenvalue larger than the mean eigenvalue. These steps were conducted in IBM SPSS Statistics software (v29.0.2.0). We estimated the mean of the scores for each principal component for each CHC trait for each population, and then estimated the pairwise Euclidean distances between populations in R v4.3.2 (66). Correlations were evaluated for significance using Mantel tests between the distances in color-pattern and in CHC traits. Mantel tests were performed in the *vegan* R package v2.6-4 (67), based on 10,000 permutations.

### Genetic covariance between color-pattern and CHCs

We estimated the genetic covariance between color-pattern and each of the CHCs traits within a population (FHA population, 34.518 N, −119.801 W). This population was chosen due to its large sample size, enabling the genetic mapping of color-pattern and CHCs traits, which was not feasible in other populations (42, 47). We note that previous studies have shown that the genetic basis for color-pattern is conserved across *T. cristinae* populations (43). We use the results of the genome wide analysis (GWA) mapping of color-pattern from (44) (https://zenodo.org/records/11050621), and performed a new GWA mapping for the different CHC traits (*i.e.,* pentacosanes, heptacosanes and nonacosanes) using the same sample set. Because color-pattern is only mapped in green and striped individuals (*i.e.,* both sexes but excluding melanistic individuals) and CHCs were only mapped in females (*i.e.,* all morphs but no males), the sample sizes were different between the two sets of data (pattern n = 538, CHC n = 197), which only partially overlapped (n for both traits = 183).

We did the GWA mapping for each CHC trait using the Bayesian sparse linear mixed models (BSLMM) in *gemma* (Markov chain Monte Carlo, MCMC, for each trait: 10 chains, sampling steps: 1,000,000; burn-in: 200,000; minor allele frequency threshold: 0 (68)). We next estimated the breeding values (BVs) based on the model-averaged effect estimates for each SNP, which includes the possible sparse/main effect and polygenic effect of each SNP. We use BVs because it is a more appropriate approach to deal with polygenic traits, characterized by small and uncertain individual effects (68, 69). We calculated genetic correlations of the BVs across traits to compute the standardized genetic covariance matrix (*i.e.,* the stantardized G-matrix). Confidence intervals on the Pearson correlations were calculated using bootstrap re-sampling of the individuals (1000 replicates each). The heritability values (the diagonal in the genetic correlation matrices) were taken as proportion of variance explained (PVE) from *gemma* outputs.

### Environmental covariation

To estimate correlations between environmental variables affecting color-pattern and CHCs, we used data from (41) (https://doi.org/10.5061/dryad.v1q13). This dataset comprises n=206 populations, n=98 on *Adenostoma* and n=108 on *Ceanothus* host plants, with information on elevation and 19 data layers from WorldClim (53). We performed Wilcoxon signed-rank tests between all climatic variables and host plants (Table S3) and among elevation and host plants. We performed Pearson correlations between climate and elevation. We did so using every climate layer with the exception of ‘precipitation of driest month’, which was invariant (*i.e.,* zero) for all populations. Additionally, we performed PCAs to summarize the data, and conducted the correlation analysis with elevation using the principal components. The first two principal components summarizing the bioclimatic data corresponded to 92.8% of the variation explained (PC1=67.9%, PC2=24.9%). The variables that contribute the most to PC1 are the precipitation variables, added to other temperature variables such as annual mean temperature (see Table S3).

We further used climate and elevation data to estimate the association between these variables and female CHCs traits. To do so, we used the CHC traits data from (47) and summarized the different traits according to their chain length (*i.e.,* pentacosanes, heptacosanes, or nonacosanes) using PCA in the IBM SPSS Statistics software (v29.0.2.0), as described above. To obtain more robust results we used data from 15 populations whose localities were also present in the dataset containing climate/elevation information (Table S2). We estimated the relationship between the first principal component axis describing CHCs variation (PC1 = 50.3%, 88.5%, and 92.0% of the variance explained for pentacosanes, heptacosanes and nonacosanes, respectively) and the first principal component axis describing climatic variation, as well as elevation (see Figs. S1-S2). All the statistical analyses were performed in R v4.3.2 (66).

### Sexual isolation

To estimate the regression relationship between divergence in each CHC trait and sexual isolation, we used the pairwise Euclidean CHC distances calculated above and the distances in sexual isolation between the 21 population pairs derived from the seven populations. We used the data from Riesch *et al.* (47), who calculated the pairwise index of sexual isolation (*I_PSI_*) based on mating propensity derived from no-choice mating trials from (70). While previous research has assessed the correlation between female CHCs and sexual isolation by aggregating all CHC traits (47), we here estimate distances for each CHC trait separately. We fitted Bayesian linear mixed models (BLMM) to estimate the degree of association between CHC divergence and sexual isolation, including random effects accounting for the pairwise nature of the variables (71, 72). The Bayesian approach uses a Markov chain Monte Carlo framework to estimate the regression coefficients and deviance information criterion (DIC) for model selection. The model was fitted via the *rjags* R package (73), including linear models (y ∼ x), and quadratic models (y ∼ x + x^2^), where divergence in each CHC class was the explanatory variable and sexual isolation was the response variable. The variables were scaled and centered before the analyses. We ran three chains of the model, with 10,000 iterations, a burn-in of 2,000 iterations, and a thinning interval of 5. The results are all represented in Table S4. Additionally, we assessed the combined effects of the CHC traits on sexual isolation. We also included geographical distances in the model, which were calculated using the geodesic distance between coordinate points and then logarithmically transformed (74). We ran BLMM using the same parameters described above. The results are represented in Table S5. All the statistical analyses were again performed in R v4.3.2 (66).

## Acknowledgments

This study is part of a project that has received funding from the European Research Council (ERC) to PN, under the European Union’s Horizon 2020 research and innovation programme (Grant agreement No. 770826 EE-Dynamics). We thank Vanina Tonzo, Laura Zamorano, Etsuko Nonaka and Henry Truchassout for insightful discussions. PN and CFdC were supported by Royal Society of London (RG140369) and FAPESP 2020/07556-8.

## Notes

### Competing Interest Statement

The authors have declared no competing interest.

