## Supplementary Information for "Ecology not genetics explains correlated trait divergence during speciation"

**This PDF file includes:**

Supporting text

Figures S1 to S3

Tables S1 to S5

Supplementary Materials

Supporting text

*Genome-wide genetic differentiation between* Timema *species*

To illustrate patterns of genetic differentiation representative of different species in *Timema*, we estimated patterns of genome-wide genetic differentiation between two *Timema* species that co-occur in nature: *T. californicum* and *T. poppensis* (47). To this end, we based on previously published whole genome sequence, using allele frequency estimates from sympatric populations of *T. californicum* and *T. poppensis* (locality LP, see Supplementary Table 3 from (47) for 5,018,138 SNPs (filtered from a set of 5,074,942 SNPs to retain only SNPs with data for at least five individuals per species). The allele frequency estimates were taken directly from this past study and were based on an earlier, more fragmented genome assembly for a melanic *T. cristinae* with scaffolds combined into linkage groups based on inheritance patterns in mapping families (see (41)).To quantify genetic differentiation, we estimated F_ST_ in 100 SNP windows as F_ST_ = Σ_i_(H_T_-H_S_)/Σ_i_(H_T_), where H_S_ and H_T_ are the mean subpopulation expected heterozygosity and the total expected heterozygosity given the mean allele frequencies, respectively. This analysis was conducted in R (66).

*Patterns of CHC variation within and among* Timema *species*

We conducted new analyses to summarize patterns of CHC variation within and among *Timema* species based on previously published data (52). The data were obtained from Dryad (<https://doi.org/10.5061/dryad.98f8c>) and comprised relative abundances of seven cuticular hydrocarbons of the heptacosane class for 76 individuals from nine *Timema* species (stick insects were sampled from two populations in most species). Using R, we performed a PCA ordination of five of the CHCs (two of the CHCs were excluded as they were not scored in three species, *i.e.,* were missing data): 13Me27, 7Me27, 5Me27, 3Me27, 9Me27+11Me27 (these are all 27 carbon molecules with methyl (Me) groups on different carbons). For this analysis, we centered and standardized the CHC data. The first two PCs explained 54.2% and 21.9% of the CHC variation, respectively. 90% data ellipses for PC scores for each species were computed using the *ordiellipse* function in the R package *vegan* (75).

Supplementary Figures

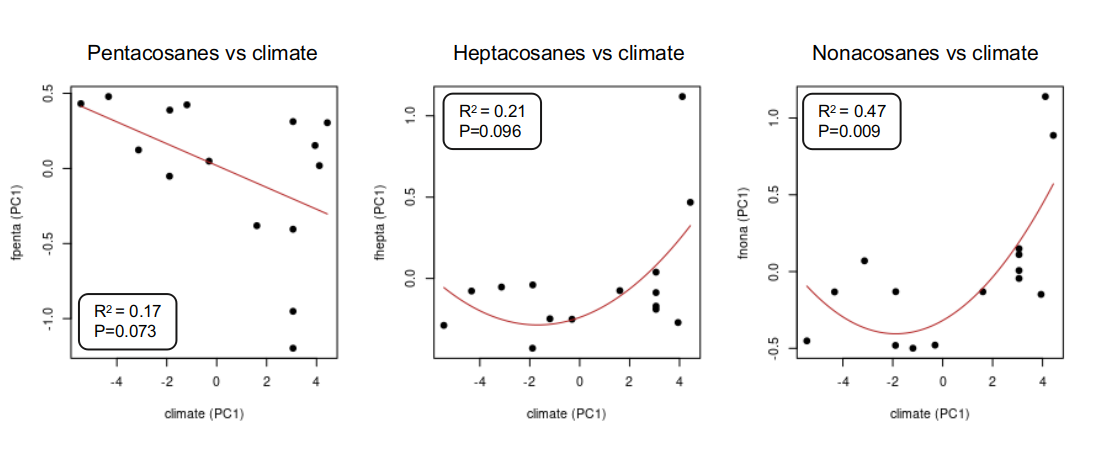
**Figure S1. Association between female CHCs traits and climatic variables**. CHCs traits from 15 populations (Table S2) were summarized for each CHC class using a principal component analysis (PCA). Here, we used the first PC axis (representing 50.3%, 88.5%, and 92.0% of the variation in female pentacosanes, heptacosanes, and nonacosanes, respectively). We used the first PC summarizing the 19 WorldClim bioclimatic variables to represent climatic variation (PC1 represents 67.9% of the variation). We used linear models to describe the regression, with the adjusted R^2^ and corresponding p-value represented in each graph. Abbreviations: fpenta = female pentacosanes; fhepta = female heptacosanes; fnona = female nonacosanes.

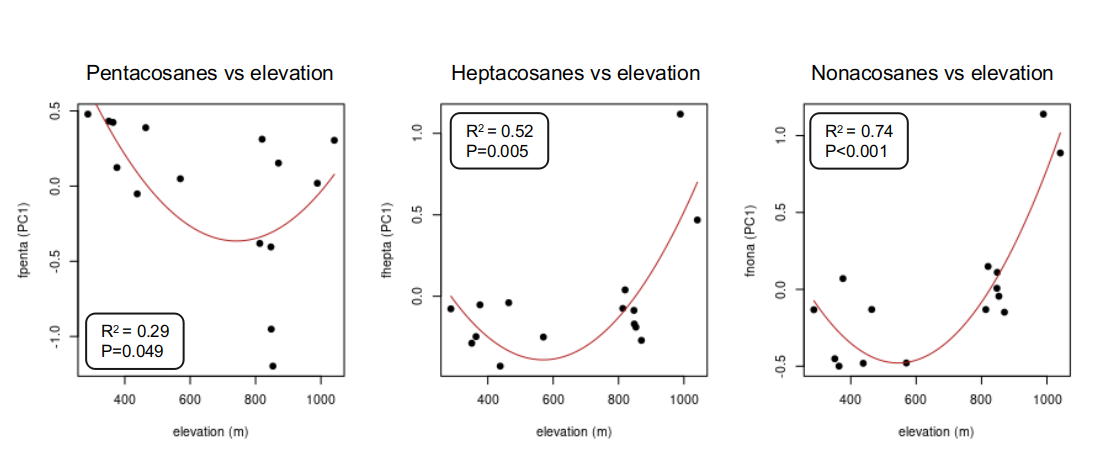

**Figure S2. Association between female CHCs traits and elevation**. CHCs traits from 15 populations (Table S2) were summarized for each CHC class using a principal component analysis (PCA), as in Fig. S1. We used linear models to describe the regression, with the adjusted R^2^ and corresponding p-value represented in each graph. Abbreviations: fpenta = female pentacosanes; fhepta = female heptacosanes; fnona = female nonacosanes.

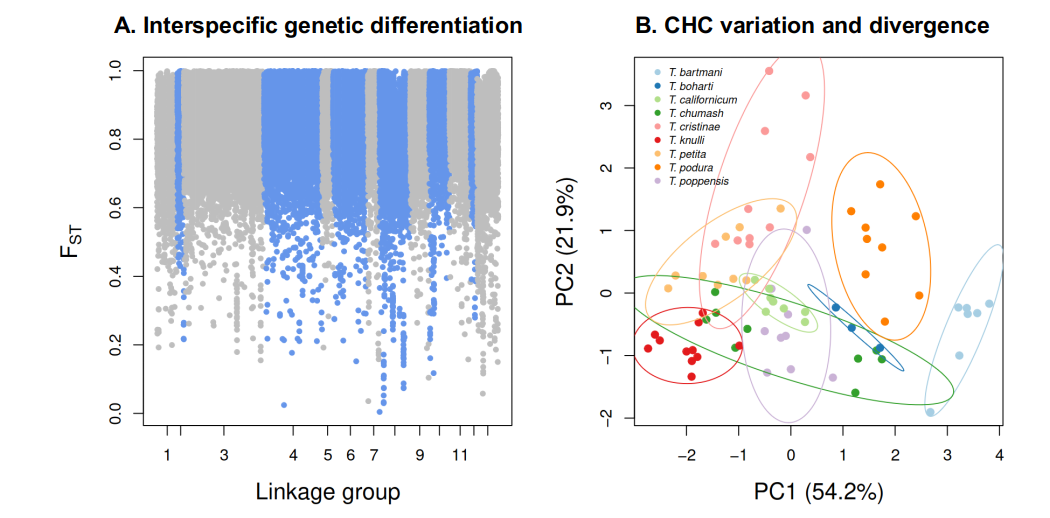
**Figure S3. Species differences among *Timema.*** (A) Manhattan plot depicting 100 SNP window estimates of F_ST_ between sympatric *Timema* species, *T. califonricum* and *T. poppensis*, from whole genome sequence data (data from (47)). (B) Principal component analysis ordination of CHC variation within and among *Timema* species. Each point represents a stick insect and is colored by species. 90% data ellipses are shown for each species (data from (52)).

Supplementary Tables

Table S1. The seven *T. cristinae* populations used to estimate divergence in color-pattern and in CHCs.

| **population code** | **locality** | **host** | **latitude** | **longitude** | **elevation (m)** |
| --- | --- | --- | --- | --- | --- |
| LA | Laurel Springs | *Adenostoma* | 34.509 | -119.796 | 820 |
| PRC | Paradise Road | *Ceanothus* | 34.533 | -119.857 | 364 |
| MA | Mattress | *Adenostoma* | 34.514 | -119.800 | 848 |
| PC | Poppy | *Ceanothus* | 34.480 | -119.770 | 287 |
| OUTA | Outlook | *Adenostoma* | 34.530 | -119.840 | 464 |
| OGC | Open Grass | *Ceanothus* | 34.510 | -119.800 | 847 |
| HVA | Hidden Valley | *Adenostoma* | 34.488 | -119.787 | 376 |

**Table S2.** *Timema cristinae* populations used to estimate the association between female CHCs and climate and elevation.

| **population** | **locality** | **host** | **latitude** | **longitude** | **elevation (m)** |
| --- | --- | --- | --- | --- | --- |
| BYA | Brick yard | A | 34.469 | -119.677 | 870 |
| ECC35A | East Camino Cielo 35 | A | 34.506 | -119.768 | 1041 |
| ECCCampA | East Camino Cielo Camp | A | 34.506 | -119.762 | 989 |
| FHA | Far Hill | A | 34.518 | -119.801 | 813 |
| HVA | Hidden Valley | A | 34.488 | -119.787 | 376 |
| LA | Laurel Springs | A | 34.509 | -119.796 | 820 |
| MA | Mattress | A | 34.513 | -119.796 | 848 |
| OGA | Open Grass | A | 34.513 | -119.796 | 853 |
| OGC | Open Grass | C | 34.532 | -119.843 | 847 |
| OUTA | Outlook | A | 34.477 | -119.769 | 464 |
| PC | Poppy | C | 34.477 | -119.769 | 287 |
| PRC | Paradise Road | C | 34.533 | -119.857 | 364 |
| R12C | Refugio 12 | C | 34.515 | -120.071 | 351 |
| R23A | Refugio 23 | A | 34.518 | -120.077 | 438 |
| SC | Stage Coach | C | 34.523 | -119.832 | 570 |

Abbreviations: A = *Adenostoma fasciculatum*; C = *Ceanothus spinosus*

**Table S3. WorldClim bioclimatic variables, contribution to the different axes of the principal component analysis, and correlation with host plant.** Correlations were estimated with Wilcoxon signed-rank test, and the table represents the corresponding *W-value* and *p-value*. The layer 14 (precipitation of the driest month) was excluded because it was zero across all localities. All correlations are significant, with the exception of between ‘maximum temperature of warmest month’ and host plant.

| **Worldclim bioclim layer** | **PC** | **W** | **p-value** |
| --- | --- | --- | --- |
| Annual mean temperature | PC1 | 2686 | 1.39e-09 |
| Mean diurnal range | PC2 | 3741.5 | 3.26e-04 |
| Isothermality | PC1 | 3044.5 | 1.61e-07 |
| Temperature seasonality | PC1 | 7431 | 2.38e-07 |
| Max temperature of warmest month | PC2 | 4911 | 4.33e-01 |
| Min temperature of coldest month | PC1 | 2800 | 7.58e-09 |
| Temperature annual range | PC2 | 7008.5 | 2.82e-05 |
| Mean temperature of wettest quarter | PC1 | 2682.5 | 1.43e-09 |
| Mean temperature of driest quarter | PC2 | 3309 | 4.75e-06 |
| Mean temperature of warmest quarter | PC2 | 3287.5 | 3.72e-06 |
| Mean temperature of coldest quarter | PC1 | 2726 | 2.62e-09 |
| Annual precipitation | PC1 | 7899 | 3.54e-10 |
| Precipitation of wettest month | PC1 | 7274 | 1.34e-06 |
| Precipitation seasonality | PC2 | 3090.5 | 2.12e-07 |
| Precipitation of wettest quarter | PC1 | 7617.5 | 1.96e-08 |
| Precipitation of driest quarter | PC1 | 7026 | 8.84e-07 |
| Precipitation of warmest quarter | PC1 | 7757 | 1.84e-09 |
| Precipitation of coldest quarter | PC1 | 7641.5 | 1.44e-08 |

**Table S4.** Summary of model comparison in Bayesian regressions evaluating the possible effects of different female CHC classes on sexual isolation (ipsi). Here, each class is evaluated alone. Among them, we evaluate pentacosanes (fpenta), heptacosanes (fpenta) and nonacosanes (fnona). We test the linear and quadratic models to explain sexual isolation. Values represented here are the deviance information criterion (DIC), and the best models are on top highlighted in bold.

| **model (fpenta vs ipsi)** | **Deviance** | **pD** | **DIC** | **ΔDIC** |
| --- | --- | --- | --- | --- |
| fpenta+fpenta² | 38.73 | 6.59 | **45.32** | 0 |
| fpenta | 43.63 | 5.48 | 49.10 | 3.78 |
| **model (fnona vs ipsi)** | **Deviance** | **pD** | **DIC** | **ΔDIC** |
| fhepta | 60.83 | 3.95 | **64.77** | 0 |
| fhepta+fhepta² | 61.55 | 5.09 | 66.64 | 1.87 |
| **model (fnona vs ipsi)** | **Deviance** | **pD** | **DIC** | **ΔDIC** |
| fnona | 61.10 | 3.96 | **65.06** | 0.0 |
| fnona+fnona² | 61.05 | 5.05 | 66.10 | 1.04 |

**Table S5.** Summary of model comparison in Bayesian regressions evaluating the possible effects of different female CHC classes on sexual isolation, namely: pentacosanes (Fpenta), heptacosanes (Fpenta) and nonacosanes (Fnona), and geographical distance (Geo). Values represented here are the deviance information criterion (DIC), supporting Fpenta alone as the best model (β_FPENTA_= 0.28 [-0.18, 0.72; 95% ETPI]). Here, we did not consider the quadratic relationship between the CHC classes and sexual isolation.

| **model** | **Deviance** | **pD** | **DIC** | **ΔDIC** |
| --- | --- | --- | --- | --- |
| **Fpent** | 59.73 | 3.82 | 63.54 | 0.00 |
| **Geo** | 59.52 | 4.21 | 63.73 | 0.19 |
| **Geo+Fpent** | 58.99 | 4.94 | 63.93 | 0.39 |
| **Fnona** | 61.00 | 3.89 | 64.90 | 1.36 |
| **Geo+Fpent+Fnona** | 59.18 | 5.93 | 65.11 | 1.57 |
| **Fhept** | 61.12 | 4.00 | 65.12 | 1.58 |
| **Geo+Fnona** | 60.07 | 5.34 | 65.42 | 1.88 |
| **Fpent+Fnona** | 60.65 | 4.79 | 65.43 | 1.89 |
| **Fpent+Fhepta** | 60.76 | 4.85 | 65.62 | 2.08 |
| **Geo+Fhept** | 60.36 | 5.41 | 65.77 | 2.23 |
| **Geo+Fpent+Fhept** | 59.87 | 6.06 | 65.93 | 2.39 |
| **Fhept+Fnona** | 62.05 | 4.93 | 66.98 | 3.44 |
| **Geo+Fhept+Fnona** | 61.05 | 6.29 | 67.34 | 3.80 |
| **Fpent+Fhept+Fnona** | 61.67 | 5.95 | 67.62 | 4.08 |
| **Geo+Fpent+Fhept+Fnona** | 60.39 | 7.25 | 67.65 | 4.11 |
